# No baby boom of social mammals in European zoos resulting from COVID-19 lockdowns

**DOI:** 10.1101/2025.03.05.641480

**Authors:** Audrey Maille, Cécile Garcia, Fernando Colchero, Jean-François Lemaître, Samuel Pavard

## Abstract

Most zoo animals are exposed daily to human visitors, which could be a major disturbance on reproduction. Although few studies have examined the relationship between number of births and crowd size by comparing high and low visitor days, the effect of zoo visitors on animal reproduction over longer timescales has been overlooked. We assessed whether number of births in 34 social mammal species hosted in zoos from 23 European countries differed after prolonged periods of closure to the public (COVID-19 lockdowns) as compared with preceding years. An analysis restricted to the periods following the lockdowns and corresponding to any conception during closure (from 2020 to 2022), compared with the same periods in the two previous years (2018 and 2019), showed no effect of zoo closure on number of births, even when considering each species’ life history traits and whether they were managed under a breeding programme. We found no evidence that a complete and prolonged absence of visitors affected the ability to reproduce in social mammals housed across European zoos. By analysing birth variation in a large sample of mammalian species, this study contributes to a better knowledge of the natality of mammals in captive settings opened to the public.

## 1. Introduction

The main goals of modern zoos are to maximize the welfare (nutrition, environment, health, behaviour and mental state) of captive animals and to ensure their successful reproduction, particularly in the case of endangered species, through reproductive management (ex-situ conservation [1]). However, although most of the species in zoos are non-domesticated, they are constantly exposed to humans, at least during opening hours to the public. This exposure to humans could be a major disturbance, increasing stress and potentially affecting crucial biological functions, such as feeding and reproduction [2,3]. During the COVID-19 outbreak, the lockdowns implemented in response to the pandemic forced zoos to close their doors for prolonged and repetitive periods between March 2020 and February 2022, depending on the countries. Suspicions of a “baby boom” quickly emerged in the zoo community and the media, which was directly linked to the disappearance of visitors (*e*.*g*. [4]), although no scientific evidence has ever supported this assumption.

The physiological and behavioural responses of zoo animals to visitors have been extensively evaluated [5]. Some studies showed increased rates of glucocorticoids (a measure of physiological stress), aggressions and repetitive behaviours associated with higher numbers of visitors (*e*.*g*. [2,6–8]), but many studies also showed a neutral or positive response (*e*.*g*. [9,10]). Differences in responses of zoo animals to visitors are likely to depend on both the species and individual characteristics of the animals [11], and on environmental variables such as enclosure design, spatial proximity to the public [12,13] and climatic conditions [14]. However, contrasting results may also be related to the variety of proxies used to measure visitor effects (crowd size, daily numbers of visitors, visitor activities or noise [10,15]), and there could also be threshold effects in the number of visitors [16].

Few studies have examined the effect of visitor presence on the reproduction of zoo animals. Most of them have focused on the timing of births in mammals by testing the *“weekend effect”* hypothesis, which proposes that captive animals are more likely to give birth during periods of low visitor exposure (weekdays). None of them found evidence for a weekend effect on birth rates, whether for Cetartiodactyla, Perissodactyla, Carnivora [3] and Primates [3,17,18]. Although these studies are relevant for understanding whether visitors might influence reproductive events in zoo mammals, they only provide insight into variation in the timing of births rather than in the offspring produced during and over the years. They also focused on the effect of visitors on birthing behaviour, but neglected mating behaviour. To our knowledge, only one study showed more socio-sexual behaviour in a captive group of macaques (*Macaca silenus)* during closing time [19].

Only a handful of studies managed to observe animals that were either exposed or concealed to human visitors [20,21], because hiding animals from the public is rather unusual in zoo settings. An alternative option to accurately evaluate the visitor effect is to compare prolonged periods of time when zoos are closed or open to the public, such as in winter compared to other seasons [22]. The unprecedented situation of COVID-19 lockdowns provided some researchers with a unique opportunity to test the effect of a complete absence of visitors, at different times of the year, on behaviour of zoo animals. Apart from variations in social, feeding and comfort behaviour during the first months of closure, no change in sexual behaviour was found between the opening and closure periods [23-27], except for one group of olive baboons (*Papio papio*) that displayed more sexual interactions during lockdown [26].

In the present study, we analysed the effect of prolonged zoo closure on 34 mammalian species. Our aim was to test whether the absence of visitors had an effect on the number of births, possibly as a result of a change in mating behaviour. Since we only expected an effect of the complete and prolonged absence of visitors for species whose males and females used to live together in captive settings and were more likely to reproduce without much human intervention (e.g. transfer between enclosures, monitoring when placed in the same enclosure), we selected species that live permanently in groups composed of adults of both sexes (*i*.*e*. social mammals). We therefore focused on species whose social organization fell into the following three categories: multi-male/multi-female, one-male/multi-female and pair. We also considered whether the species were managed under breeding programmes as, for these species, recommendations may be made to temporarily suspend reproduction for the entire captive population or to prevent the breeding of some individuals in certain zoos, which could override any effect of visitors on their ability to reproduce [28]. These potential biases linked to husbandry constraints can, to some extent, be offset by a substantial amount of data and by studying a large number of species in a wide variety of zoos.

Data used in this study originate from the Zoological Information Management System (ZIMS), a real-time and centralized database of animals under human care (including information from over 1,200 zoos worldwide and records for over 22,000 species) managed by the international non-profit Species360 [29]. We expected an increased number of births after periods of lockdowns as compared with preceding years, which could potentially reflect a positive effect of the absence of visitors on reproduction of social mammals in zoos. Moreover, given that social species such as primates are more likely to give birth during the night [30], plausibly a time when the environmental and social disturbance is lowest, we predicted diurnal species to be more affected by visitor absence during daytime. In addition, we assumed that species managed under breeding programmes would be less affected by the absence of visitors, as their ability to reproduce highly depends on recommendations of studbook holders. We further investigated for any effect of the social organisation, mating system, seasonality of reproduction, gestation length and absence or presence of polytocy (production of several youngs per litter) as these traits are likely to affect reproductive behaviour of mammals throughout the year [31,32].

## 2. Methods

### (a) Data

To identify dates of closures in European zoos, we downloaded data on response policies to COVID-19 in European countries provided from the European Centre for Disease Prevention and Control (version 2022-08-25; [33]). This database includes the dates of various responses categorized into 28 modalities – from restriction on mass gatherings to stay-at-home order – for 30 European countries in a period starting on 2020-02-27 and ending on 2022-02-15. We identified two modalities that secure the closure of zoos: ‘*StayHomeOrder*’ (enforced stay-at-home orders for the general population; and also referred to as ‘lockdown’) and ‘*ClosPubAny*’ (closure of public spaces of any kind); 27 countries implemented one of these two restrictions at least once.

Data on mammal births recorded between 2010-01-01 and 2022-02-14 in European zoos were provided by Species360 and the Zoological Information Management System (ZIMS). We did not have access to the identity of zoos where the births occurred, but only to the countries where they were located. We obtained birth data for zoos in 23 of the 27 European countries for which we had data on COVID-19 responses (see **Figure 1**). We restricted our analyses on births to 34 social mammal species that are abundant in European zoos (to avoid problems due to small sample size) and for which at least 40 births were recorded in 2018-2019 (so more than 20 per year, a sample size per species that allows tests to compare the number of births between years). The 34 selected species represent a wide range of mammalian orders (5 Artiodactyla, 9 Carnivora, 2 Diprodontia, 13 Primates and 5 Rodentia; see **Table 1**).

**Table 1.** Taxonomic order, scientific and common species name, social organisation (multi-male/multi-female, one-male/multi-female, pair) and management under a breeding programme (Yes if EAZA ex-situ programme in January 2020) for the 34 mammalian species.

| Order | Species | Common name | Social organisation | Breeding programme |
| --- | --- | --- | --- | --- |
| Artiodactyla | *Antidorcas marsupialis* | Springbok | Multi-male/multi-female | Yes |
|  | *Antilope cervicapra* | Blackbuck | One-male/multi-female | No |
|  | *Madoqua kirkii* | Kirk's dik-dik | Pair | Yes |
|  | *Pecari tajacu* | Collared peccary | Multi-male/multi-female | No |
|  | *Potamochoerus porcus* | Red river hog | One-male/multi-female | Yes |
| Carnivora | *Canis lupus* | Gray wolf | Multi-male/multi-female | Yes |
|  | *Cynictis penicillata* | Yellow mongoose | One-male/multi-female | No |
|  | *Helogale parvula* | Dwarf mongoose | Multi-male/multi-female | No |
|  | *Lycaon pictus* | African wild dog | Multi-male/multi-female | Yes |
|  | *Mungos mungo* | Banded mongoose | Multi-male/multi-female | No |
|  | *Panthera leo* | Lion | Multi-male/multi-female | Yes* |
|  | *Speothos venaticus* | Bush dog | One-male/multi-female | Yes |
|  | *Suricata suricatta* | Meerkat | Multi-male/multi-female | No |
|  | *Vulpes zerda* | Fennec | Pair | Yes |
| Diprotodontia | *Macropus giganteus* | Eastern gray kangaroo | Multi-male/multi-female | Yes |
|  | *Macropus rufus* | Red kangaroo | One-male/multi-female | Yes |
| Primates | *Callimico goeldii* | Goeldi's monkey | Pair | Yes* |
|  | *Callithrix jacchus* | White-tufted-ear marmoset | Pair | Yes* |
|  | *Cebuella pygmaea* | Pygmy marmoset | Pair | Yes* |
|  | *Colobus guereza* | Guereza | Multi-male/multi-female | Yes* |
|  | *Lemur catta* | Ring-tailed lemur | Multi-male/multi-female | Yes* |
|  | *Papio hamadryas* | Hamadryas baboon | Multi-male/multi-female | Yes* |
|  | *Saguinus imperator* | Emperor tamarin | Pair | Yes* |
|  | *Saguinus midas* | Midas tamarin | Pair | Yes* |
|  | *Saguinus oedipus* | Cotton-top tamarin | Pair | Yes* |
|  | *Saimiri boliviensis* | Bolivian squirrel monkey | Multi-male/multi-female | Yes |
|  | *Saimiri sciureus* | South American squirrel monkey | Multi-male/multi-female | Yes |
|  | *Theropithecus gelada* | Gelada baboon | Multi-male/multi-female | Yes* |
|  | *Varecia variegata* | Ruffed lemur | Pair | Yes* |
| Rodentia | *Ctenodactylus gundi* | Gundi | Pair | Yes |
|  | *Cynomys ludovicianus* | Black-tailed prairie dog | Multi-male/multi-female | No |
|  | *Dolichotis patagonum* | Patagonian mara | Pair | No |
|  | *Hydrochoerus hydrochaeris* | Capybara | Multi-male/multi-female | No |
|  | *Hystrix indica* | Indian crested porcupine | Pair | No |

\*re-established after 1st January 2018 following the implementation of the new European Association of Zoo and Aquaria (EAZA) Population Management structure (https://www.eaza.net/assets/Uploads/CCC/January-2020.pdf)

**Figure 1.**
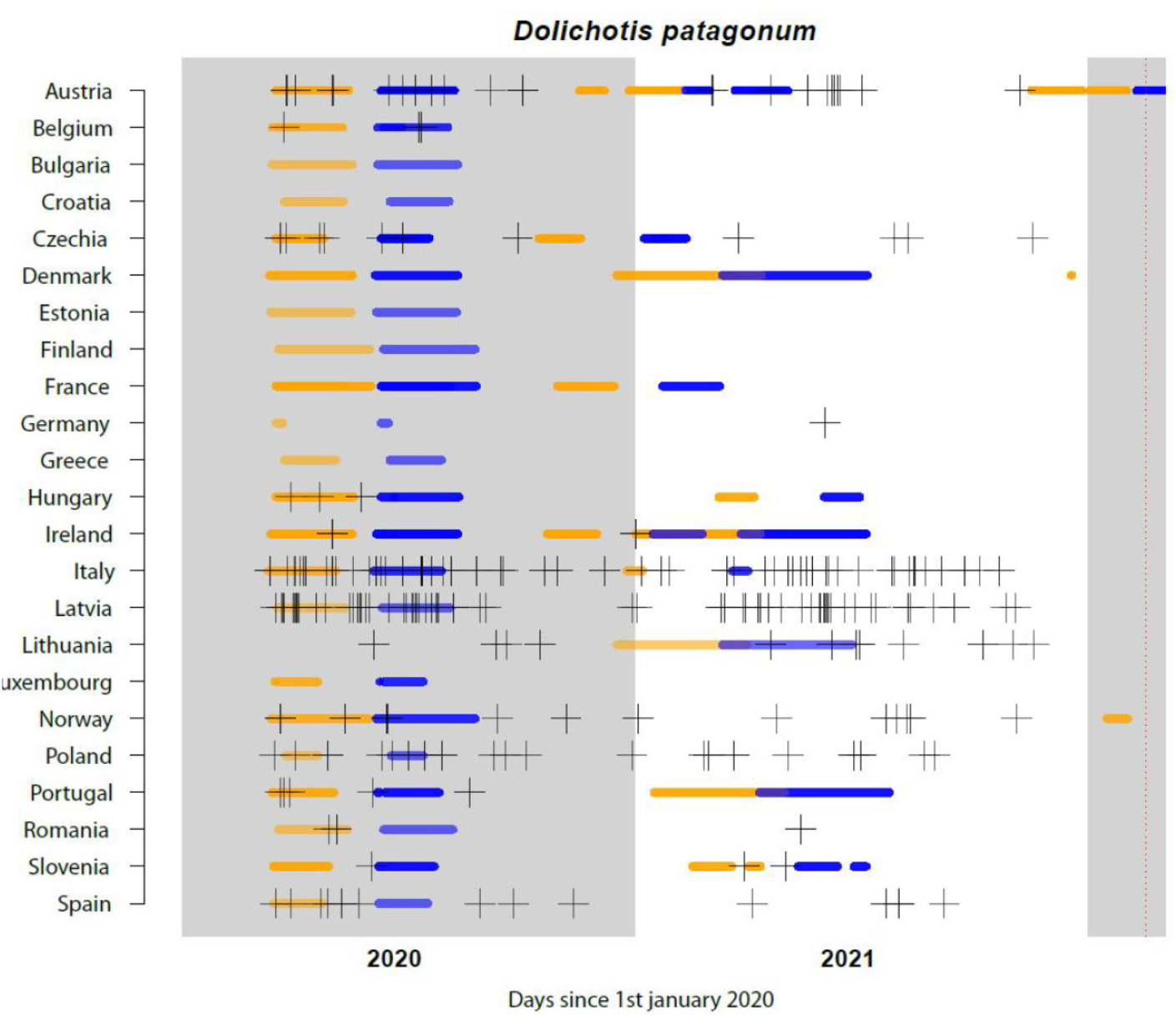
Time windows corresponding to lockdowns (in orange) and births observation (if conception occurred during lockdown, in blue) across countries in 2020 (grey) and 2021 (white), in the case of the Patagonian mara (*Dolichotis patagonum*). The crosses correspond to the dates of observed births for each country line. The Patagonian Mara was chosen because it provides several illustrative points for Austria, which experienced 5 lockdowns. As the gestation period of the Patagonian mara is 85 days, the first births observation window in 2020 is after the lockdown period. However, in 2021, successive periods of lockdown overlap with windows of births observation. The last observation period is truncated at the last date of birth observation on 14 February 2022 (vertical red dotted line).

The dataset encompasses information on date and country of birth for 31 309 individuals from those 34 species from 2010-01-01 to 2022-02-14. Although we did not have access to the births after mid-February 2022, the data provided were sufficient to capture any effect of the closures, as the last lockdown took place before May 2021 for all countries except Austria (blue lines in **Figure 1** and **Figure S1**).

For each species, the following life history traits were extracted from various online database such as Global Biodiversity Information Facility (gbif.org), Animal Diversity web (animaldiversity.org) or Encyclopedia Of Life (eol.org): circadian rhythm, social organisation, mating system, seasonality of reproduction, gestation length and polytocy (see details in **Table S1**). In regard to social organisation, 17 of the 34 species were classified as multi-male/multi-female, 12 as forming pairs and 5 as one-male/multi-female (**Table 1)**. Most species (N=24) are managed under breeding programmes coordinated by the European Association of Zoo and Aquaria (*i*.*e*. EEP for EAZA Ex-situ Programme; **Table 1**).

### (b) Analyses

We started with a global descriptive analysis, comparing the total number of births in the years of closures for COVID-19 (2020 and 2021) with the previous 10 years (2010 to 2019). We calculated the percentages of change from year to year and the average percentage of change. We then examined whether each percentage was higher than the average plus two times the standard error. The exploration of the total number of births across a large number of comparative years would reveal any deviations from the usual variation in the species considered. We assume that any fluctuation in number of births would reflect a change in fertility, which might be due to lockdowns. However, we stressed that year-to-year variations in the number of births could also be due to changes in number of zoos holding the species and number of fertile females over the years.

In a second analysis, in order to control for major changes in population composition, we compared two consecutive time frames at a shorter scale. We restricted the analysis to periods of births corresponding to any conception during lockdowns, which we called the *“births observation”* time window. We compared number of births in “births observation” time windows in 2020 and 2021 to the same periods but in 2018 and 2019. These ‘births observation’ time windows are each specific of the country (which differ in closure times) and species (which differ in gestation length). More precisely, we denote [t_1,j_, t_2,j_] the duration time of lockdowns in country *j*. Therefore, for a given species *i*, having gestation length *g*_i,_, the “births observation” time window was [t_1,j_ + *g*_i_, t_2,j_ + *g*_i_] (see examples in **Figure 1** and **Figure S1**). Number of births recorded during “births observation” time windows for each species are provided in **Table S1**. Because the last observed birth in our database occurred at t_max_= 2022-02-14, we considered only “births observation” time windows starting before this date such that t_1,j_ + *g*_i ≤_ t_max_ (therefore, time windows closed at t_max_ if t_2,j_ + *g*_i ≥_ t_max,;_ **Figures 1 and S1**). Given that the first lockdown date was March 10^th^, 2020 in Italy and the shortest gestation time was of 30 days for the black-tailed prairie dog (*Cynomys ludovicianus*), the earliest t_1,j_+g_i_ was April 9^th^, 2020, while the latest t_2,j_ + *g*_i_ was t_max_. We thus compared the total number of births per species occurring during “births observation” time windows over the period [2020-04-10, 2022-02-14] with the total number of births occurring over the same time windows in the period [2018-04- 10, 2020-02-14]), using Wilcoxon tests.

Finally, we investigated whether the species’ circadian rhythm, social organisation, mating system, seasonality of reproduction and management under a breeding programme explained the proportion of change in number of births between the two periods (*i*.*e*., [# births _[2020-04-10, 2022-02-14]_ - # births _[2018-04-10, 2020-02-14]_] / # births _[2018-04-10, 2020-02-14]_). We log-normalized this proportion (Shapiro test > 0.05) and investigated the effect of the covariates on this response variable, by using linear regressions. We first performed simple analyses and then multiple regressions also including gestation length (continuous variable) and polytocy (Boolean variable) as covariates. Indeed, species with long gestation were less likely to be affected by short or successive lockdowns. In contrast, polytocous species were expected to impact more our analysis because we analysed number of births as number of pups rather than as breeding events (*i*.*e*. the available dataset included the number of births by species and by country, without specifying the zoo where they have occurred). Model selection was based on Akaike’s Information Criterion corrected for small sample size [33]. All analyses were performed in ‘R’ version 4.3.3.

## 3. Results

The year 2020 had the largest number of annual births recorded from 2010 to 2021, with an increase of 3.7% in 2020 with respect to 2019 and 5.4% when compared to the average number of births over the previous 10 years (**Figure 2**). However, this substantial increase was followed by a considerable reduction in 2021, as the number of births were 9 % lower in 2021 compared to 2020.

**Figure 2.**
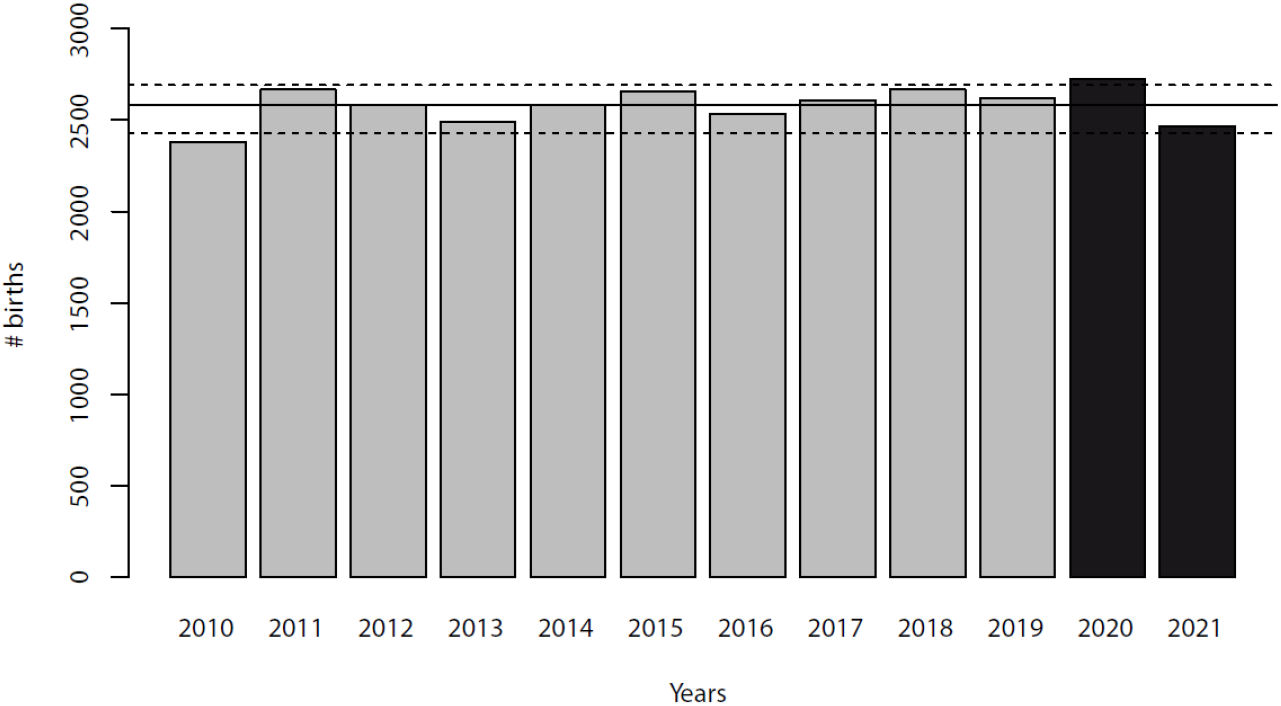
Number of births per year from 2010 to 2021, all the 34 species included.

Only 16 species had a number of births in 2020 larger than the average number of births in 2010-2019. Of these species, only 5 species had a larger number of births than average plus two times the Standard Error (**Figure S2**): the Indian crested porcupine (*Hystrix indica*: +86%), Kirk’s dik-dik (*Madoqua kirkii*: +64%), grey wolf (*Canis lupus*: +34%), Goeldi’s monkey (*Callimico goeldii*: +33%) and capybara (*Hydrochoerus hydrochaeris*: +26%).

There were 628 births in total during the “births observation” time windows from 2020 to 2022 (**Table S1**), which fell well within the observed range of values recorded in the same time windows from 2010 to 2019 (mean = 660, SD=56). A comparison between the number of births per species falling within the “births observation” time windows (from 2020 to 2022) to the number of births that have occurred within the same time windows two years before (2018 and 2019) did not show any difference between the two paired series of values (Wilcoxon: p>0.1; **Figure 3**).

**Figure 3.**
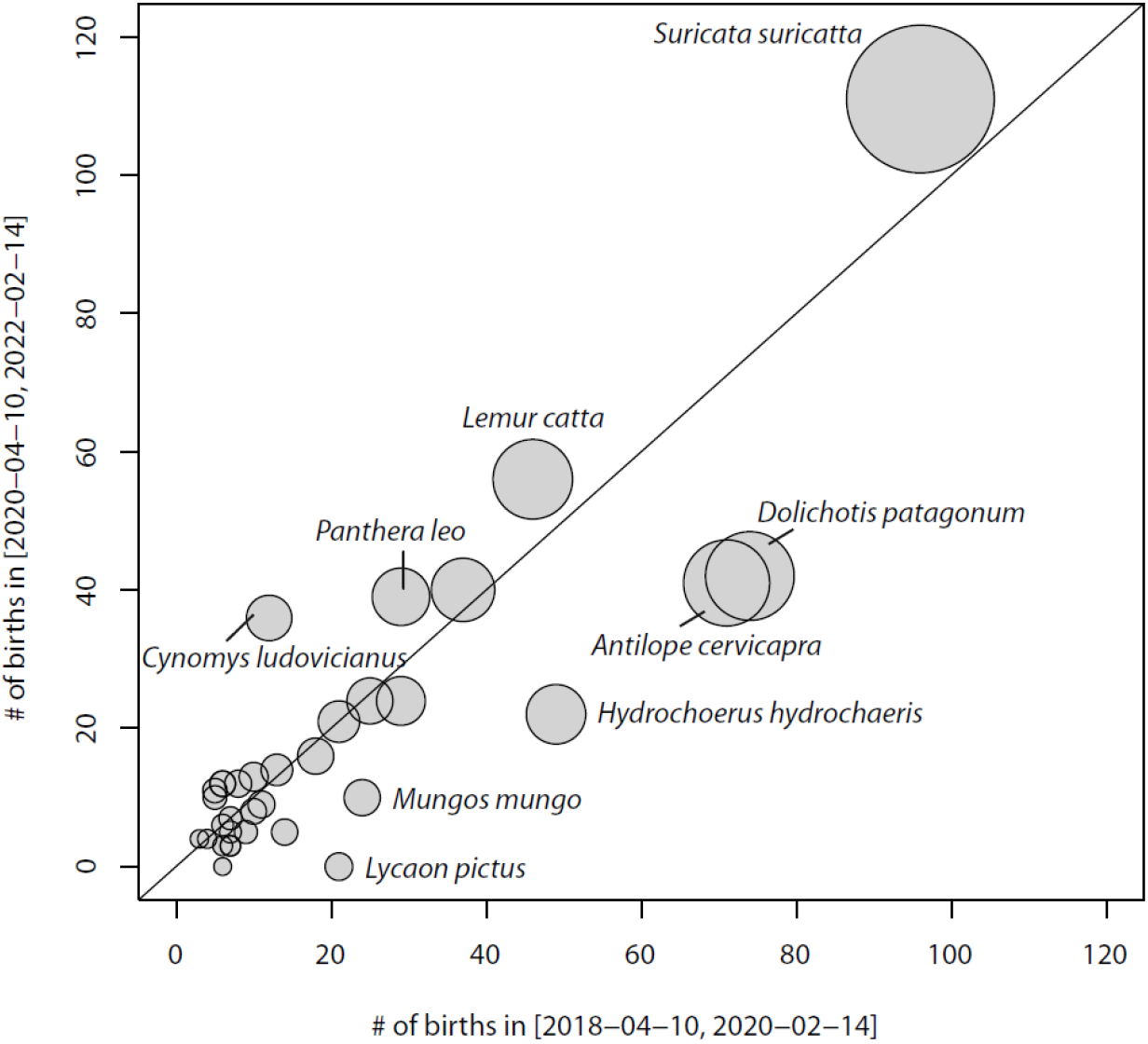
Number of births for each of the 34 species within the two periods [2020-04-10, 2022-02-14] and [2018-04-10, 2020-02-14]. Size of the points is proportional to the total number of births recorded over the two periods.

Finally, linear regressions showed no significant effect (p>0.05) of circadian rhythm, social organisation, mating system, seasonality of reproduction and implementation of a breeding programme on the proportion of change in the number of births in period [2020-04- 10, 2022-02-14] compared to [2018-04-10, 2020-02-14] (**Table 2**), whether gestation length and polytocy were included in the analyses. None of the multifactorial regressions performed better (ΔAICc>2) than the monofactorial regressions in explaining variation in number of births.

**Table 2.** Monofactorial regression of the percentage of change in the number of births falling into the *births observation* time window in period [2020-04-10, 2022-02-14] compared to period [2018-04-10, 2020-02-14] according to explanatory variables, for the 34 species. Multifactorial analyses with gestation length and polytocy as covariates give similar results.

|  |  | Estimate | SE | t-value | Pr(>\|t\|) |
| --- | --- | --- | --- | --- | --- |
| **Circadian rhythm** | Diurnal (n=23) |  |  |  |  |
|  | Cathemeral (n=3) | -0.05 | 0.19 | -0.30 | 0.77 |
|  | Nocturnal (n=8) | -0.14 | 0.13 | -1.06 | 0.30 |
| **Social organisation** | Multi-male (n=17) |  |  |  |  |
|  | One-male (n=5) | -0.16 | 0.16 | -1.02 | 0.32 |
|  | Pair (n=12) | 0.1 | 0.12 | 0.82 | 0.42 |
| **Mating system†** | Monogamous (n=12) |  |  |  |  |
|  | Polyandrous (n=5) | 0.32 | 0.16 | 2.04 | 0.053 |
|  | Polygynous (n=9) | -0.02 | 0.13 | -0.18 | 0.86 |
|  | Promiscuous (n=8) | 0.15 | 0.14 | 1.12 | 0.27 |
| **Seasonal reproduction** | No (n=12) |  |  |  |  |
|  | Yes (n=21) | -0.05 | 0.11 | -0.45 | 0.66 |
| **Breeding programme** | No (n=10) |  |  |  |  |
|  | Yes (n=24) | 0.05 | 0.12 | 0.43 | 0.69 |
| **Gestation length (yrs)** |  | 0.24 | 0.41 | 0.60 | 0.55 |
| **Polytocy** | Polytocous (n=23) |  |  |  |  |
|  | Monotocous (n=11) | -0.016 | 0.12 | -0.14 | 0.89 |

† Regrouping together polyandrous and promiscuous species give qualitatively similar results

## 4. Discussion

We showed that the complete and prolonged absence of visitors, during COVID-19 lockdowns, did not lead to an improvement in natality in 34 social mammal species housed in zoos from 23 European countries. This lack of effect is unlikely caused by a decrease in the number of reproductive females over time, as we analysed the number of births both at a large scale (annual variation over 12 years) and at a short scale (comparison of periods corresponding to any conception during lockdown with the same periods but two years earlier).

Descriptive analyses do not show an increase in the number of births, for all species combined, in 2020 compared to the following year (2021) and the previous ten years (2010 to 2019). Of the 34 species, five species experienced an increase in birth number in 2020 followed by a decrease in 2021, including one Artiodactyla, one Carnivora, one Primate, and two Rodentia. They might have paid the cost of an increased allocation to reproduction in 2020, resulting in a lower number of births in the following year. Costs of reproduction on subsequent reproduction are expected to particularly affect species displaying long gestation length [35] and polytocy [36]), but these life history traits had no effect on the number of births recorded in 2020 and 2021 in the 34 sampled species. Three of the five species, Kirk’s dik-dik, grey wolf and Goeldi’s monkey, were managed under a breeding programme. Fluctuation from a period of increasing births (2020) to a period of decreasing births (2021) in those species may reflect a responsible population management following a birth surplus, with reproduction suspended in certain animals until the current offspring have been transferred to other zoos. Alternatively, as spring 2020 was exceptionally hot and dry in Western Europe [37], the species-specific peak in births recorded that year may be related to climatic factors as weather variation can affect both behaviour [14] and reproduction [32].

An analysis restricted to the periods following the lockdowns (“births observation” time windows), compared with the same periods in the two previous years, also showed no effect of prolonged zoo closure on the number of births, even when considering each species’ life-history traits and whether they were managed under a breeding programme. A performance test showed that it was unlikely that we would fail to detect a 10% increase (while there would have been a one in two chance of missing an increase of less than 5%; **Analysis S1**). Yet, those findings have several limitations. Firstly, major changes in the management of zoo animals may have counteracted the effects of lockdowns. Staff organisation was often disrupted by lockdowns, leading to a reduction in the management of animal reproduction, even for species under breeding programmes. In addition, the suspension of transfers of many animals during lockdowns (exacerbated by the implementation of Brexit in February 2020, which stopped transfers with the United Kingdom [38]) may have led to congestion or shortages of certain animals, which would have masked the effects of visitors. Secondly, a major caveat of the present study lays in the lack of fine behavioural observations (*e*.*g*. copulation rates) or physiological measurements (*e*.*g*. gestation and abortion rates) associated with birth observations, which prevents us from going further in interpreting visitor effects on reproductive physiology and reproductive output. For instance, one cannot exclude that the absence of visitors during lockdowns had a positive effect on ovulation and conception rates, even if this was not followed by an increase in the number of births [39].

In conclusion, our study finds no evidence that the prolonged absence of visitors influenced the ability to reproduce in social mammal species housed across European zoos. Future work integrating physiological and behavioural measurements would further clarify how the zoo environment can affect reproductive outputs.

## Supporting information

Supplemental Figure S1, Figure S2, Analysis S1

## Acknowledgements

This research was made possible by the worldwide information network of zoos and aquariums members of Species360. This study was authorised by the Species360 Research Data Use Agreement 83970.

