## Supplemental Figure S1, Figure S2, Analysis S1 for "No baby boom of social mammals in European zoos resulting from COVID-19 lockdowns"

**Table S1** – Summed data per country for the 34 species considered for the analyses (Excel file)

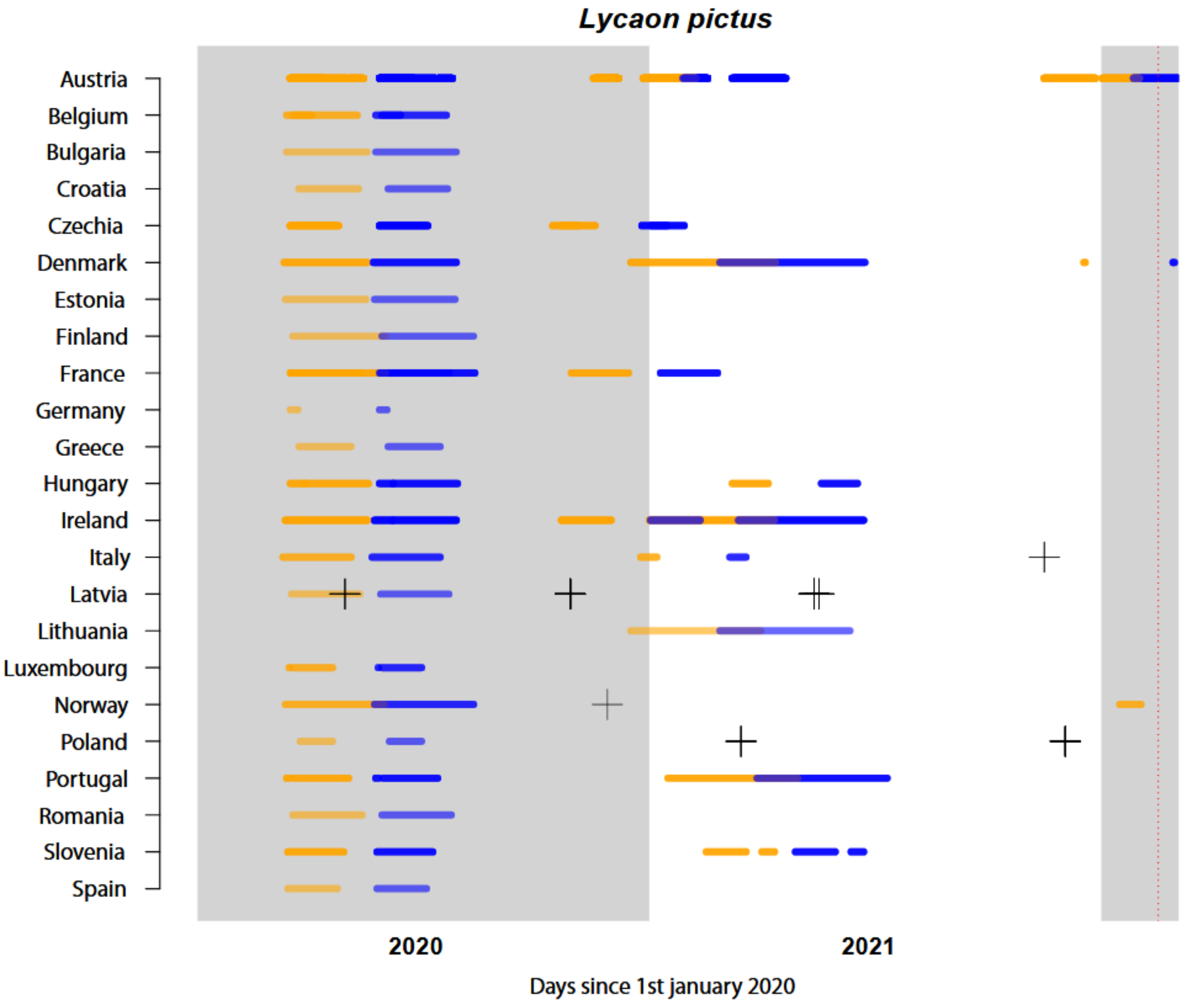


**Figure S1 –** Time windows corresponding to lockdowns (in orange) and time windows of births observations (if conception occurred during lockdown, in blue) across countries during years 2020 (grey) and 2021 (white), in the case of the *African wild dog (Lycaon pictus*), which has a gestation length of 72 days. Crosses correspond to the dates of observed births: 45 births were recorded in the period [2020-04-10, 2022-02-14] but these correspond only to 8 reproductive events (*Lycaon pictu*s having litter size about 10-11). Over these 8 reproductive events occurring in 4 different countries, none of them falls in “births observation” time windows (see also Table S1 that shows zero births were recorded in time windows in period [2020-04-10, 2022-02-14] while 21 births were recorded in time windows in period [2018-04-10, 2020-02-14]). The last observation period is truncated at the last date of birth observation on 14 February 2022 (vertical red dotted line).

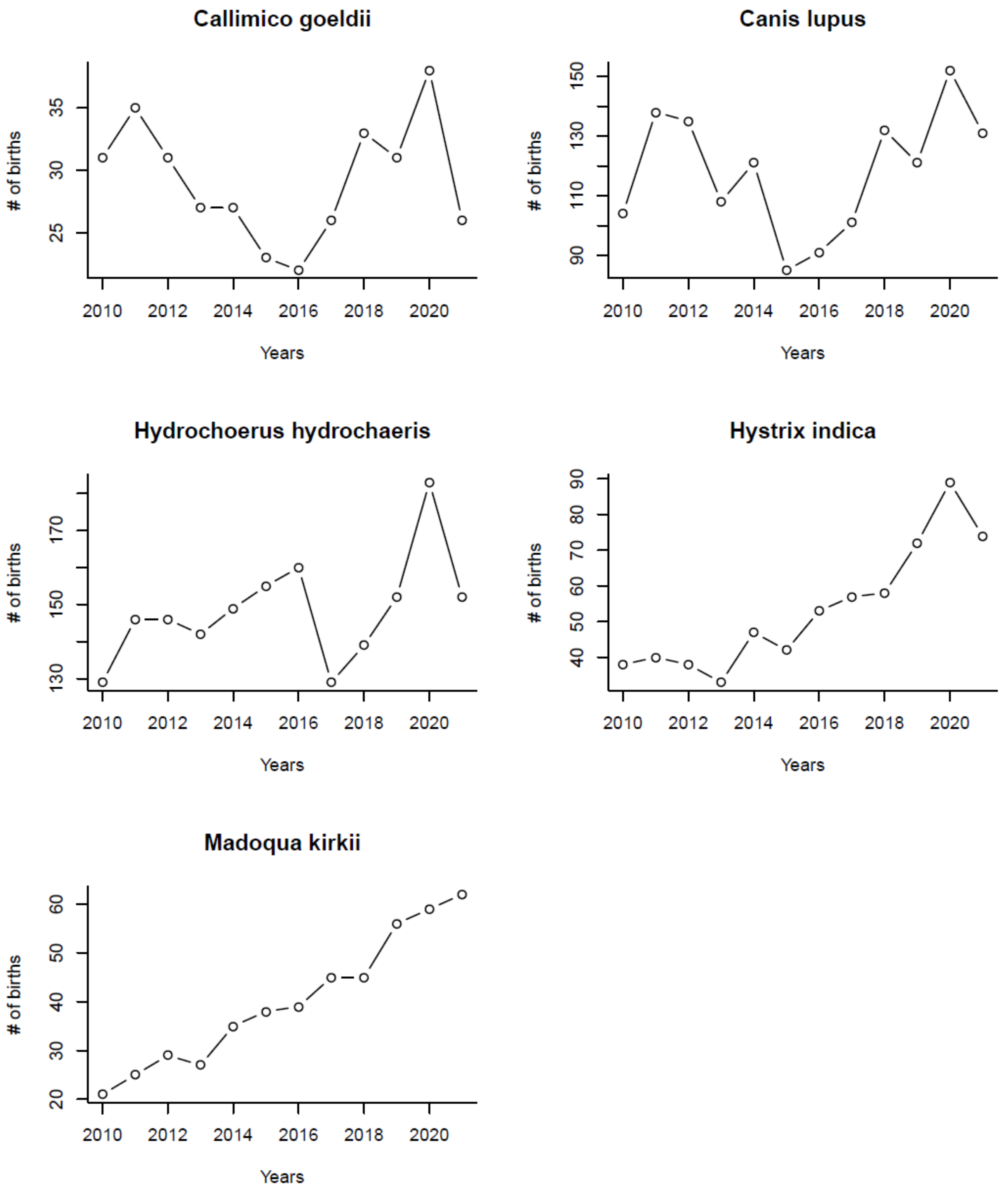


**Figure S2 – Yearly number of births from 2010 to 2021 for the five species for which values in 2021 is larger than the 2010 to 2019 average plus two times the Standard Error.** For all species, peaks in 2021 can be interpreted as a continuation of the increase in the number of births during the preceding years.

**Analysis S1** – Power test

We analysed number of births per species that occurred in time windows corresponding to any conception during lockdowns, which we called the *"births observation"* time window, and we compared them with the same periods but in the two previous years. To evaluate our ability to detect an effect, if any, a power test was performed.

Duration of each lockdown varied between countries: in country *j*, the *"lockdown"* time window was [$t_{1,j}$ , $t_{2,j}$]. Therefore, for a given species *i*, having gestation length $g_{i,}$, "births observation" time window was [$t_{1,j} + g_i$, $t_{2,j} + g_i$], which varied between countries as it depended on the duration of the lockdown in a country. Because the last observed birth in our database occurred at $t_{max}$= 2022-02-14, we considered only "births observation" time windows starting before this date such that $t_{1,j} + g_i \leq t_{max}$ (therefore, time windows closed at $t_{max}$ if $t_{2,j} + g_i \geq t_{max,;}$ **Figures 1 and S1**). Given that the first lockdown date was March 10th, 2020 in Italy and the shortest gestation time was of 30 days for the black-tailed prairie dog (*Cynomys ludovicianus*), the earliest $t_{1,j}+g_i$ was April 9th, 2020, while the latest $t_{2,j} + g_i$ was $t_{max}$. We thus compared the total number of births per species occurring during "births observation" time windows over the period [2020-04-10, 2022-02-14] with the total number of births occurring over the same time windows but two years before (*i.e.* [2018-04-10, 2020-02-14]), using Wilcoxon tests.

To test the performance of the analysis to detect any effect, we ran a power test. Let **b** and **v** be the respective vectors of per-species mean and variance in the number of births falling within the "births observation" time window from years 2010 to 2019. We determined the per-species number of births that could have been observed in period [2020-04-10, 2022-02-14] in absence of effect by adding to **b** a random number sampled in a normal distribution of mean equals to zero and variance equal to **v**. To this number was also added, for each species *i*, the number $\mathbf{b}_i k$, where *k* is the potential increased proportion of births within [2020-04-10, 2022-02-14] for species *i*. We then tested 1000 times our ability to significantly detect the increased number of births in period [2020-04-10, 2022-02-14] compared to period [2018-04-10, 2020-02-14] with $\alpha$=5%.

Results of power tests showed that we would have detected a 5% increase in number of births in about 50% of cases and a 10% increase in about 96% of cases.
